# Computation or Weight Adaptation? Rethinking the Role of Plasticity in Learning

**DOI:** 10.1101/2024.03.07.583890

**Authors:** Gili Lior, Yuval Shalev, Gabriel Stanovsky, Ariel Goldstein

## Abstract

The human brain is an adaptive learning system that can generalize to new tasks and unfamiliar environments. The traditional view is that such adaptive behavior requires a structural change of the learning system (e.g., via neural plasticity). In this work, we use artificial neural networks, specifically large language models (LLMs), to challenge this traditional view and suggest that such adaptive behavior can be also achieved solely through computation if the learning system is sufficiently trained. We focus on statistical learning paradigms. These require identifying underlying regularities in seemingly arbitrary word sequences and are primarily considered to require neural plasticity. LLMs can capture arbitrary structures without requiring weight adaptation, despite divergence from their natural language training data. Our work provides novel insights into the role of plasticity in learning, demonstrating that sufficiently trained learning systems are highly flexible, adapting to new tasks and environments solely through computation, much more than previously acknowledged. Furthermore, our work opens the door for future research to use deep learning models to conjure hypotheses about the brain.^1^

## Introduction

The human brain is a learning system that can quickly adapt to new environments and tasks using previously acquired knowledge and experiences. For example, riding a bicycle is a complex skill requiring substantial practice. However, it would not be surprising if an experienced skateboard rider could excel at bicycle riding after just a few attempts, leveraging their previously acquired balance and coordination skills. Such adaptations, often referred to as transfer learning (Ellis, 1965), are commonly viewed as forms of brain plasticity and are considered fundamental to learning new skills.

Learning in humans raises a fundamental question: Is plasticity inherently required for complex adaptive behavior, or can it be performed solely through computations without changing the system’s structure? Addressing this question has the potential to reshape our understanding of the essential components underlying adaptive behavior. Demonstrating that such behavior could emerge without structural changes might alter how we understand and study learning mechanisms in humans.

To address this question in a controlled setting, we turn to large language models (LLMs), and test their *in-context learning* (ICL) abilities on the statistical learning task, which is largely considered to require plasticity in humans (Conway, 2020). As elaborated below, ICL offers an equivalent for non-plastic learning, where a static model is shown several examples in the prompt, from which it is required to generalize to a new question. By testing artificial neural networks in ICL setups on the statistical learning task, we can shed light on whether the task inherently requires plasticity, or whether it is possible for learning systems to master it without structural changes.

In-context learning in LLMs operates entirely through activation dynamics, rather than weight updates - suggesting that learning can emerge from transient internal states (von Oswald et al., 2022; Xie et al., 2021). Interestingly, previous work has demonstrated correlations between model activations during task performance and patterns of neural activity in the human brain (Caucheteux & King, 2022; Goldstein et al., 2022, 2023, 2024; Schrimpf et al., 2021), hinting at possible functional similarities between LLM computations and cognitive processes. While such correlations do not imply identical mechanisms, they open up exciting possibilities for using LLMs as computational models of brain-like inference, particularly in tasks that require flexible, context-dependent behavior.

We test LLMs in ICL setups on the statistical learning task, as it is widely used to test *human* learning capabilities and is considered to require brain plasticity (Conway, 2020). The statistical learning paradigm measures participants’ ability to observe samples from a new environment following some arbitrary regularity (e.g., dictated by a context-free grammar) and predict outcomes consistent with its underlying structure (Saffran et al., 1996; A. Schapiro & Turk-Browne, 2015). Learning in this task occurs as the cognitive system’s behavior is adapted when exposed to new samples and structures (Aslin & Newport, 2012; Brady & Oliva, 2008). We simulate two learning tasks, inspired by two different statistical learning paradigms: Artificial Grammar Learning (AGL; (Reber, 1967)) and the Serial Reaction Time Task (SRTT; (Nissen & Bullemer, 1987)). SRTT is a motor learning task that measures reaction time to stimuli. Participants in this task show a decrease in their response times to stimuli drawn from a repeated sequence. This paradigm indicates that people can adapt their expectations to a sequence structure without knowing there is an underlying regularity in the sequence they observe (Forest et al., 2022; Robertson, 2007). The AGL paradigm is an implicit learning task where participants are presented with sequences of words that follow the rules of a regular language. Afterwards, they are asked to classify the validity of new sequences (i.e., whether they belong to that language or not). Participants in this task are not instructed to search for any regularity in the stimuli. Still, their ability to determine the validity of unseen sequences indicates that they have managed to learn the underlying regularities of the regular language. These paradigms are well established in the field of statistical learning and demonstrate the ability of humans to adapt their behavior to unrecognized statistical structures. Interestingly, these two tasks diverge from the natural language used for training LLMs. Yet, we show that LLMs can adapt to statistical patterns similar to those underlying the stimuli in these tasks without plasticity.

This work leverages LLMs to probe whether statistical learning can emerge from pure computation, formulating SRTT and AGL as in-context learning tasks. The experimental design, as described in Fig. 1, involves providing the LLM with a contextual prefix of a specific statistical regularity. We then quantify the model’s adaptation by measuring the conditional probability it assigns to valid, rule-consistent continuations. An increase in this next-token probability, as a function of the prefix length, would serve as evidence for computational, in-context adaptation to the underlying statistical structure.

**Figure 1:**
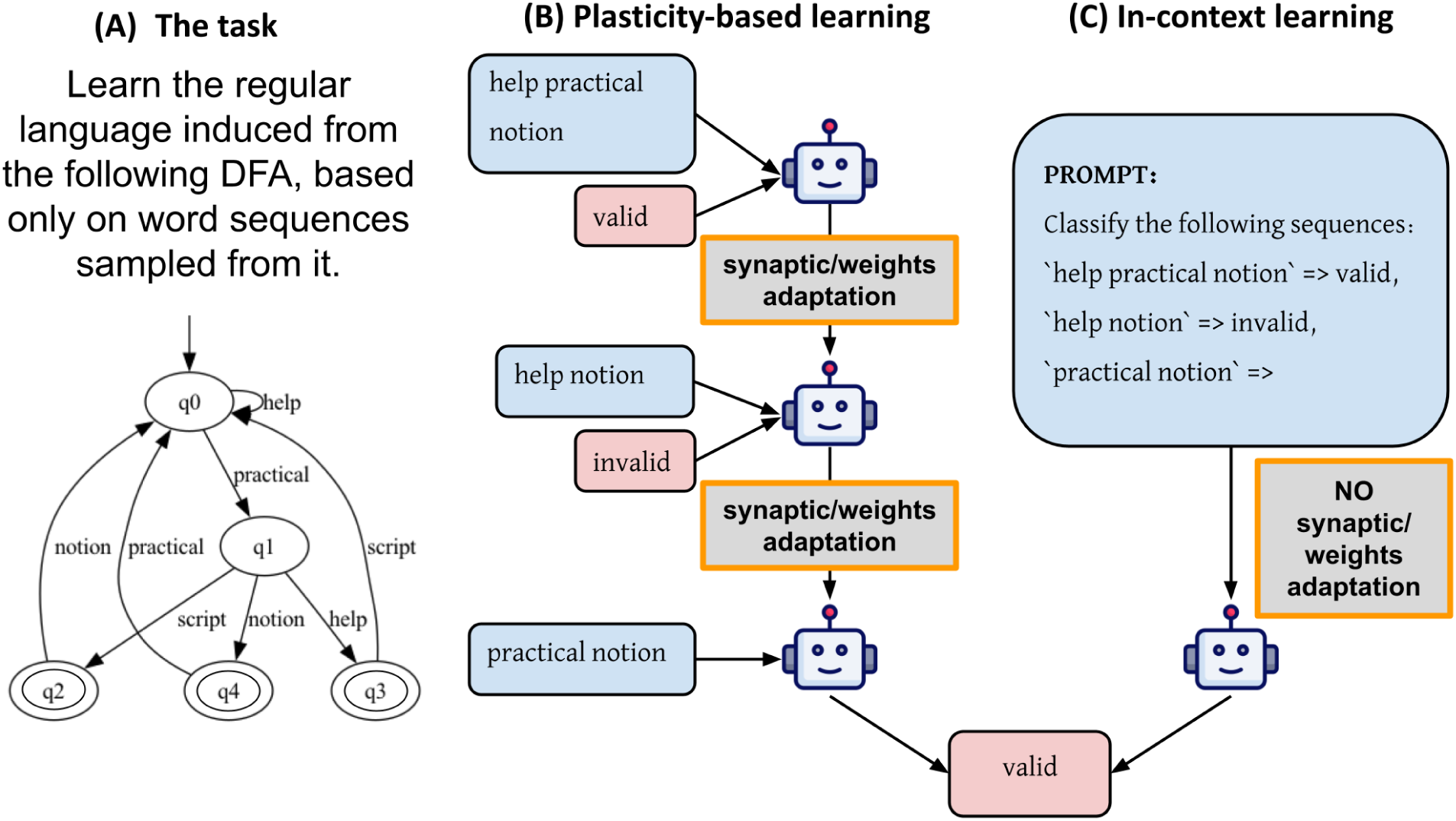
Demonstrating plasticity-based vs in-context learning. (A) Example for statistical learning task to identify a regular artificial language induced from a DFA. It can be learned via (B) plasticity-based learning, i.e., fine-tuning, or (C) In-context learning (ICL) paradigms. Fine-tuning involves plasticity, as the model updates its parameters following feedback over its predictions (gradient update), while ICL only involves prompting with a few demonstrations of the task.

We find compelling evidence that both SRTT and AGL patterns are learnable via computation, as the model’s predictions rapidly converge on the true probabilities of the sequence-generating grammar within just a few hundred tokens of exposure. Crucially, this adaptation is not limited to improving the probability of the single correct next token. Instead, the LLM appears to model the underlying state transitions of the grammar. This is evident in the AGL task where, for any state with multiple possible continuations, the model simultaneously elevates the probabilities for all valid next tokens. This reflects that the model adapts to the entire statistical structure rather than just a single sequential association.

Our findings suggest a broader spectrum of potential learning mechanisms that could coexist or complement synaptic plasticity’s role in human cognition. Furthermore, this work showcases how advancements in artificial intelligence, particularly deep learning, provide novel insights into foundational concepts like learning and plasticity. It presents a new scientific perspective to the broader cognitive science community.

## Results

Our experiments are designed to test the possibility of learning structures such as the ones that are learned in statistical learning paradigms without weight adaptation. In particular, we focus on the recent Mistral model (Jiang et al., 2023) in learning arbitrary structures without changing their inner state, i.e., without updating their parameters (weights). We specifically choose the Mistral LLM as it is currently one of the best-performing open models (Jiang et al., 2023), and in the supporting information (SI) we show that our findings hold for other off-the-shelf LLMs.^2^

We formulate two statistical learning tasks to test the ability of LLMs to capture the underlying regularities of a statistical structure, which can be represented as a directed graph with transition probabilities between states (Fig. 2A). In this paradigm, learning is demonstrated when the LLM assigns significantly higher probabilities to valid next tokens than to invalid ones within sequences of the corresponding regular language. First, inspired by SRTT, we test the abilities of an LLM to capture deterministic regularities, i.e., each state is connected to a single consecutive state. In SRTT, a reduction in participants’ reaction time to stimuli reflects that the stimuli become more expected, which means that participants learn the underlying regularities of the sequence, similarly to LLMs learning the underlying regularities of a deterministic graph.

**Figure 2:**
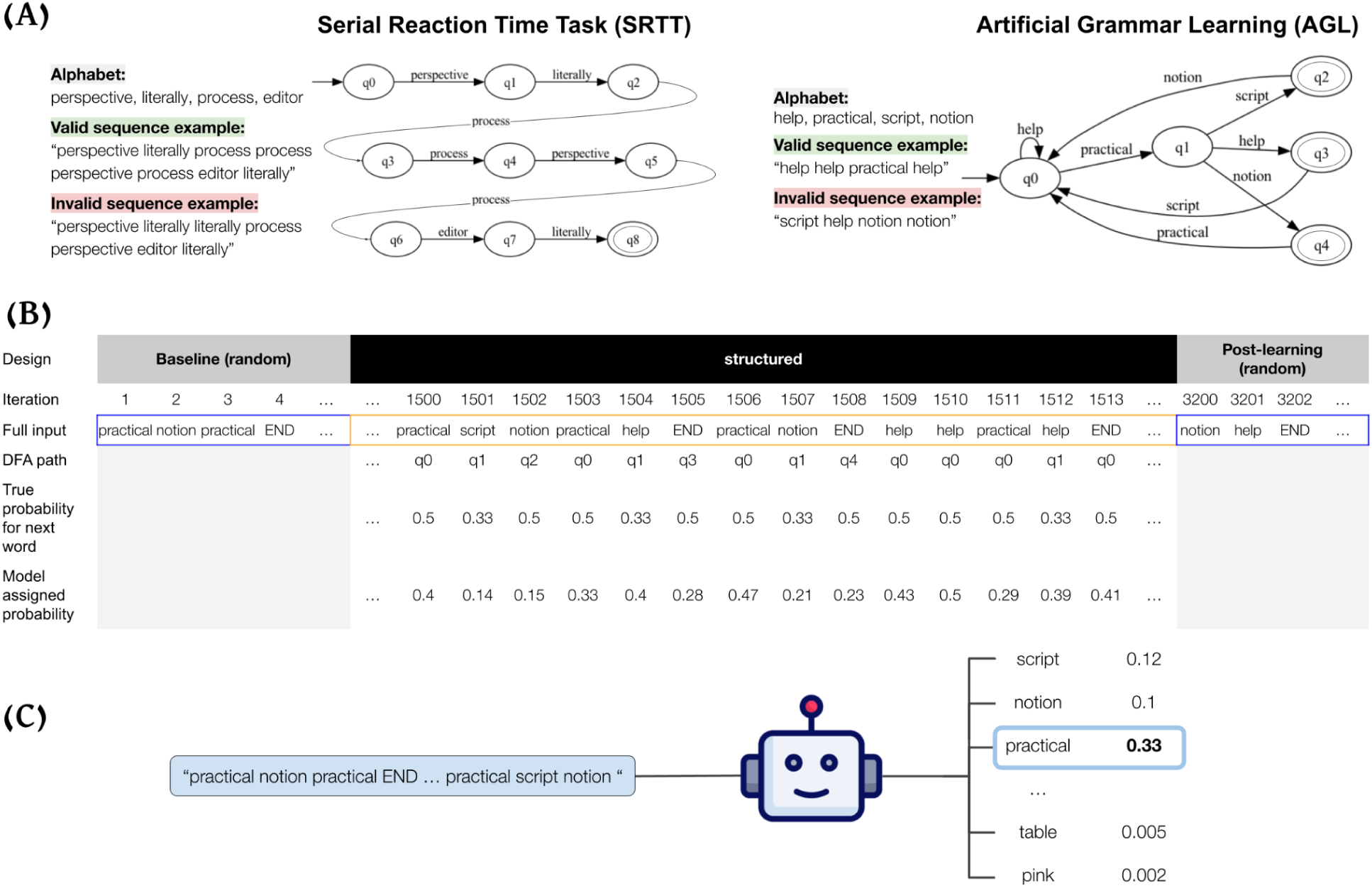
The experimental setup of our work. (A) The DFAs that simulate SRTT and AGL, with the graph alphabet and the induced valid and invalid sequences. The AGL is inspired by Pothos(Pothos, 2007). (B) An input example for artificial grammar learning consists of random sequences followed by valid sequences induced from the DFA and random sequences. This sequence is induced by traversing the DFA, following its transition probabilities, which are also presented in the table. (C) Single iteration example, where we feed the model with a prefix of our input, then the model assigns a probability for each word in its vocabulary to be the next word, and we extract the probability that the model assigns to the actual next input word. Our experiment consists of len(input) iterations, where in each iteration, we add one more word to the prefix we feed the model with.

Second, inspired by the AGL task, we test LLMs’ ability to learn non-deterministic regularities. In AGL, participants are presented with multiple sequences sampled from a regular language. The validity classification task as formulated in AGL can be simulated using a deterministic finite automaton (DFA; Fig. 2A). To test LLMs, we formulate the task as follows: given a prefix of a sequence, we test the probabilities that the model assigns to all possible tokens in some language’s vocabulary. To generate multiple sequences congruent with the AGL-like experiment, we perform a random walk on a graph that is similar to the DFA that forms the regular language, which is further explained in section A in SI, and demonstrated in Fig. S1.

We do not perform model parameter updates in our experimental setup; we only use an incremental input setup. In each iteration, we feed the model with a single additional word that is concatenated to all previous sequences of words (Fig. 2B). This new word is a valid continuation of a sequence, corresponding to each of the tasks. After each concatenation of a word, we extract the probability that the model assigns to the next word (Fig. 2C). We will show model flexibility in learning the underlying probabilities without changing its inner state, but only by feeding it with a longer input congruent with its regularity.

Inspired by classic SRTT experiments (Robertson, 2007), our experiment consists of three parts. First is a baseline part of randomly generated sequences with uniform probability over a chosen alphabet (i.e., invalid sequences). This part’s goal is to eliminate all biases of the pre-trained LLM towards specific sequences. Next is the structured part, which consists of sequences induced by traversing the corresponding task’s DFA, i.e., valid sequences. Last is the post-learning part of randomly generated sequences again (i.e., invalid sequences). In this paradigm, learning is demonstrated by an ascent in performance between the baseline part (first part) and the structured part (second part). The improvement of the performance is defined by the increase in the probability that the LLM assigns to the correct next token. Part three, which we refer to as post-learning, was designed as a control for the possibility that the LLM simply increases the probabilities of all the words from the vocabulary of the DFA.

To eliminate possible biases related to the specific vocabulary choice, our results are the mean of 100 different runs, where in each run, we choose a different vocabulary and randomly sample different sequences for both baseline and structured parts. For robustness, we report results across additional language models and different DFAs in Figs. S3–S4, as well as across alternative vocabularies—such as letters, digits, and symbols—in Fig. S10. In addition, we perform the same experiments on an untrained network, showing that the ability to adapt to such sequences arises from the pretrained model’s learned representations, rather than from its architecture alone (Fig. S11).

In Fig. 3, we illustrate the model’s predictive capabilities by presenting the probability assigned to the input’s congruent next word in each iteration in SRTT type experiment (Fig. 3A) and AGL type experiment (Fig. 3B). In both experiments, at the baseline part, the model quickly adjusts its predictions to the equally distributed transition probabilities between words in some vocabulary. Though we address this part as a baseline, it still requires LLM to learn some statistical structure without additional training. However, it is a much simpler structure - it can be thought of as a graph with a single state, with a single edge to itself, and all words in the vocabulary are of that edge label. These induced sequences are easier to learn in-context, as the model only needs to learn the vocabulary, and not transitions, which makes it a much easier task. Then in the structured part, for both SRTT (Fig. 3A) and AGL (Fig. 3B), we see the expected learning pattern - an ascent from chance probabilities at the baseline random part (depicted in blue) towards the theoretical transition probabilities upper bound at the structured part. The predictions are depicted in orange, and the upper bound is in a black dashed line; to explain how it is calculated, see section B in SI. Due to the deterministic transitions of the SRTT, the theoretical upper bound for the next token probability is 1, while for the AGL, it is an average of 0.45. The model reaches 92% of the SRTT upper bound and 79% of the AGL upper bound, averaging the last 1000 iterations of the structured part. As expected, we also see a descent back to chance probabilities at the post-learning random part (in blue). We further provide a statistical analysis of the results, presented in Fig. S5 In the SI, as well as a more granular view into the model’s learning trajectory in the AGL experiment, by examining the distribution of prediction probabilities for varying sequence lengths (Fig. S9).

**Figure 3:**
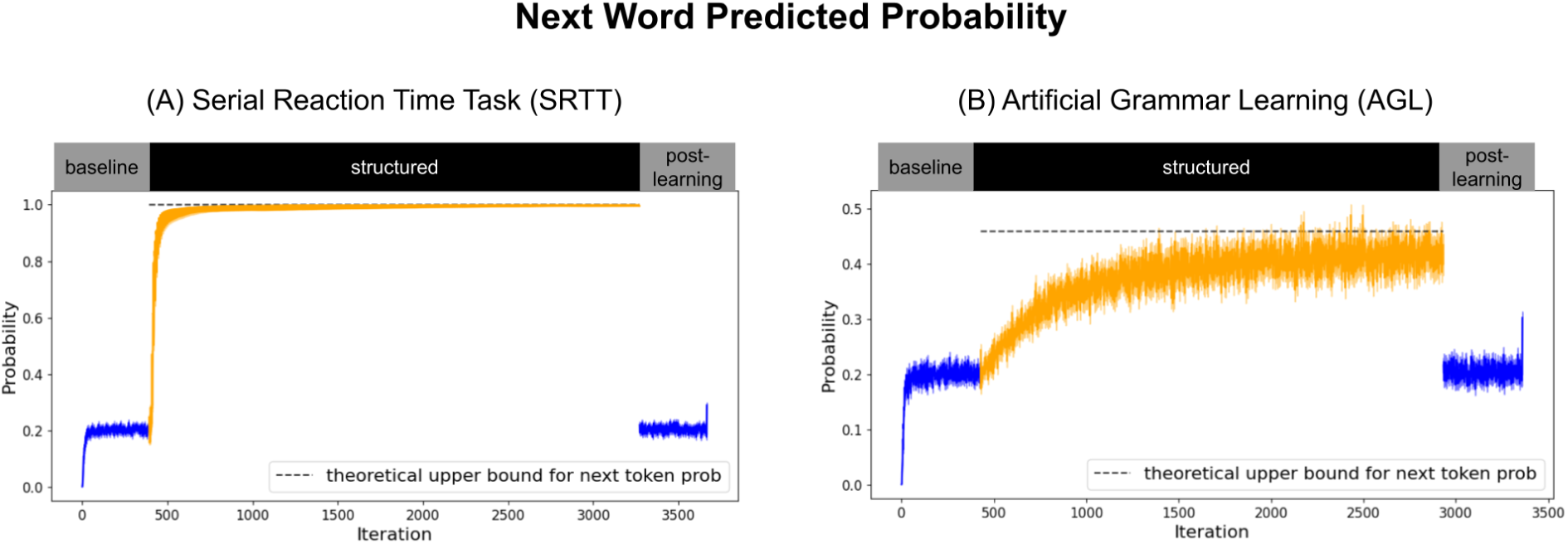
Next word probability for SRTT and AGL. Blue is the baseline part, which are random words in random sequences, and orange is the probability of words combining sequences induced by the corresponding DFA. The black dashed line is the theoretical upper bound of the next word probability, which for SRTT sequential trials is 1, as its DFA is deterministic, and for the AGL, it is ∼0.45.

We replicate the results with models having context windows smaller than 1024 tokens, as shown in Fig. S12. Specifically, we reproduce the experiment using GPT-2 (Radford et al., 2019) and find that learning emerges after fewer than 100 tokens, suggesting that a 4k-token context window is not necessary for acquiring the AGL.

To better align our setup with human AGL experiments, where participants are typically exposed to only subtle grammatical violations, we conducted another experiment, with two conditions. Both conditions share an identical initial (“training”) phase without weight adaptation but differ in the subsequent testing phase: one condition uses exclusively valid sequences, while the other introduces minimal grammatical errors by swapping two consecutive tokens. Results show that LLMs, despite no weight adaptation, assign significantly higher probabilities to valid continuations, indicating sensitivity to subtle grammatical violations comparable to human learners (Fig. 4).

**Figure 4:**
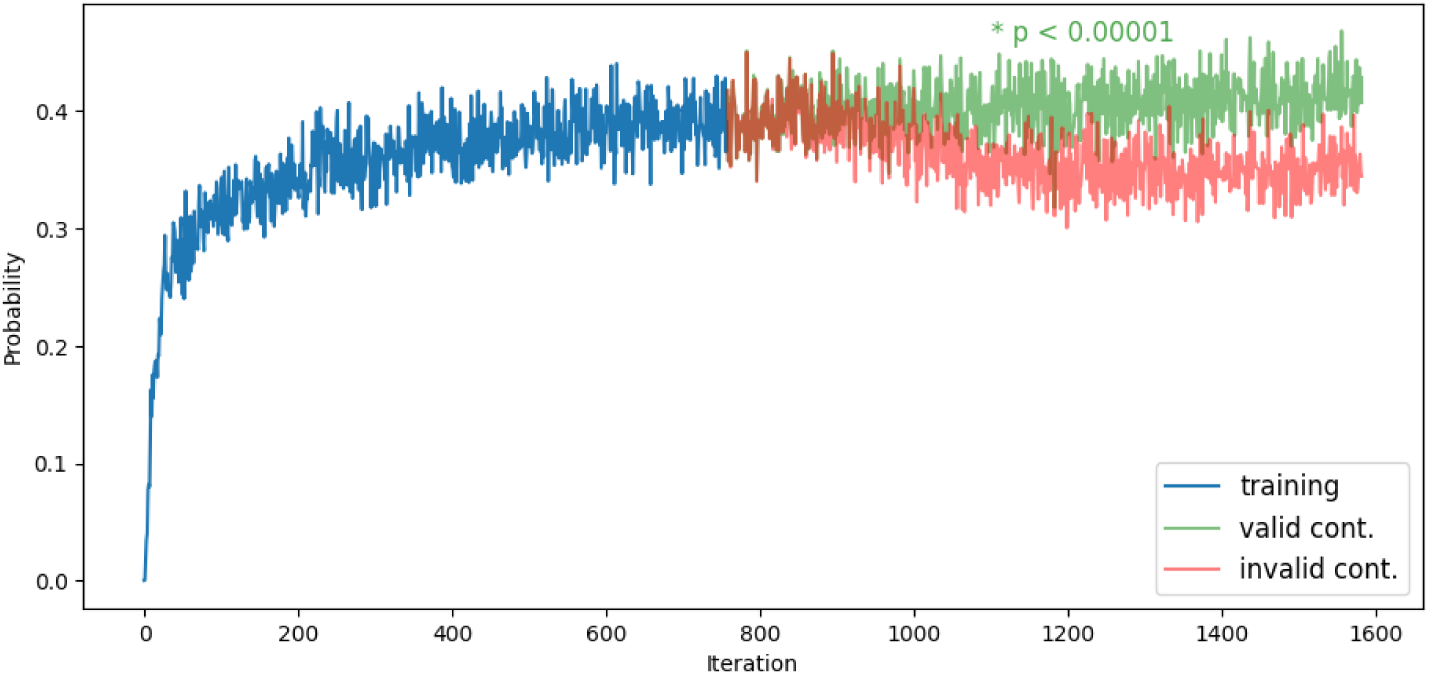
Model probability trends with minimal-error invalid sequences. The plot illustrates the probability assigned by the model to different sequence types over iterations. In the first experiment (blue and orange), the model is exposed only to valid sequences throughout. In the second experiment (blue and green), the model starts with valid sequences but transitions to minimally altered invalid sequences midway through, where two consecutive tokens in each sequence are swapped. Despite these minor grammatical errors, the model assigns significantly higher probabilities to valid continuations (p < 0.00001), demonstrating its ability to distinguish subtle violations and adapt to the correct statistical structure without weight adaptation.

One possible hypothesis for the model assigning the correct probabilities is that the model may memorize sequences and not learn the underlying structure that induces the language. To control this alternative explanation, we run an experiment where we drop all duplicating sequences in the model input (∼10% duplicate sequences). As presented in Fig. 5A, we find the learning pattern of an ascent in probabilities between the baseline and the structured part, indicating that the model does not rely on memorization but captures the underlying statistical structure. Importantly, while this manipulation removes repetition across sequences, it does not eliminate *within-sequence repetition*— that is, repeated elements (e.g., transitions or states) within a single sequence. Due to the structure of the DFA, such internal repetitions occur in approximately 75% of sequences. Thus, in this respect, the model is exposed to similar types of repetition as in human AGL studies, where limited generalization has often been linked to familiarity with recurring elements (Altmann et al., 1995; Tunney & Altmann, 2001).

**Figure 5:**
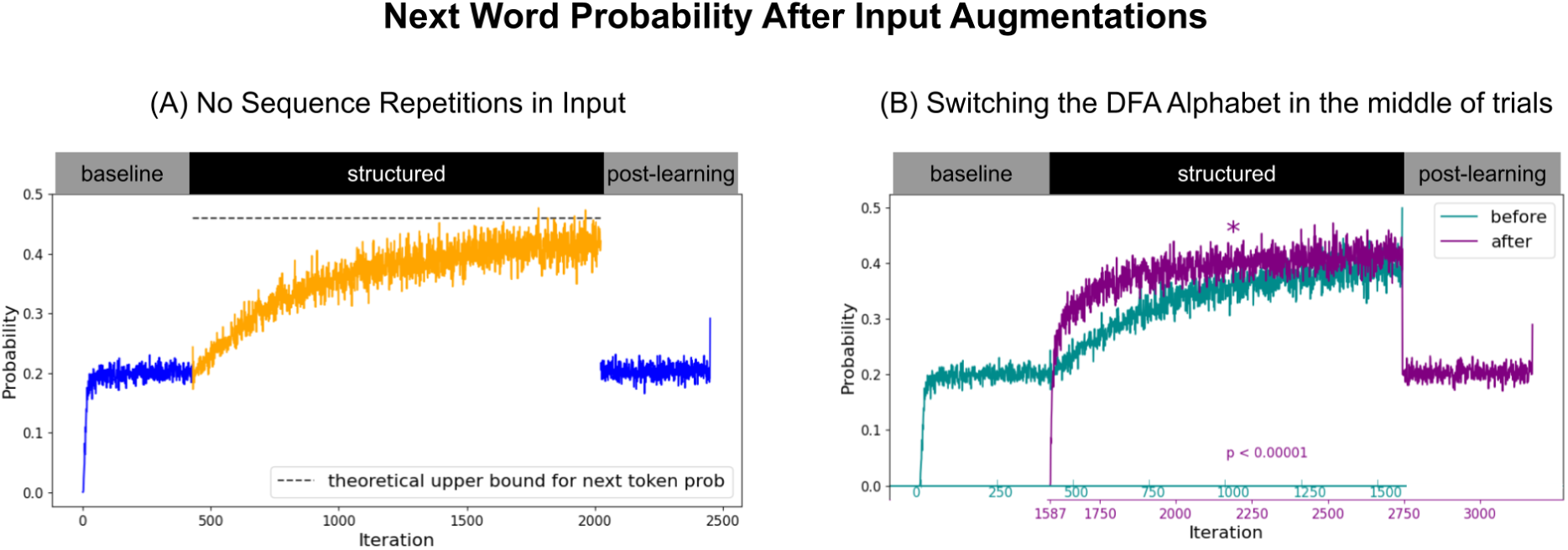
Next word probability for two AGL experiments with some input augmentation. Figure A on the left presents the probability for an input with no repeating sequences, indicating that the model does not use memorization in its predictions. On the right, we present the probability of switching the DFA alphabet in the middle of the input, which results in sequences with the same structure but consisting of different words. The purple graph, which is the sequence probability after the switch, increases quicker than at the beginning of the sequential trials (in blue), indicating that the recovery from the switch is faster than the initial learning, hence evidence for the model learning the abstract structure of the graph and not just an association between the words shown in its input.

We also propose a setup that controls for an alternative explanation: that the model may simply learn associations between specific input words rather than the underlying structure. Evidence from human studies suggests otherwise – people can transfer their knowledge of an underlying structure from one vocabulary to another, adapting rapidly to the new words (Altmann et al., 1995; Tunney & Altmann, 2001). To parallel this in our framework, the experiment in Fig. 5B switches the DFA alphabet midway through input generation (at trial 1587): the graph traversal remains unchanged, but the edge labels are replaced with new words, producing sequences with the same structure but a different vocabulary. In comparing the mean predicted probability before and after the switch, a paired t-test revealed a significant difference (t(1156)=-44.67, p=4.62e-254). This indicates that the model learned the structure that induces the sequences, rather than the specific words within them. These findings align with a previous line of work regarding humans’ ability to transfer learning in AGL (Pothos, 2007).

A potential confound for the increased learning rate after the vocabulary switch may stem from the initial exposure to random sequences, making it harder for the model to identify the structured rules of the regular language, necessitating an additional effort to discard previously acquired patterns from the unstructured, random phase. To explore this potential confound, we conduct a series of experiments varying the length of the random baseline phase. Our findings indicate that a longer random baseline correlates with slower learning of the structured phase, as depicted in Fig. S6 of the SI. However, learning rate improvements after the vocabulary switch remain significantly pronounced, even without exposure to random sequences. Fig. S7 in the SI compares the learning at the first 100 iterations at the start of the structured phase versus the first 100 iterations following the vocabulary switch, indicating a significantly increased learning after the switch. This reinforces our conclusion that even when the edge labels are changed, the model learns a statistical structure and can generalize the DFA structure.

A fundamental trait of a regular language that is induced by some DFA, is that a prefix of a sequence has several valid continuations, which is represented in the DFA with multiple outgoing edges from a certain state. A model that learns a regular language should also reflect this aspect, as demonstrated in Fig. 6. In each iteration, we extract the probabilities of all possible continuations and compare them with the graph’s transition probabilities at each state. In Fig. 6, we show the probabilities the model assigns each word given some prefix that ends at q1, and in Fig. S7 in SI, we provide a similar analysis for all other DFA states.

**Figure 6:**
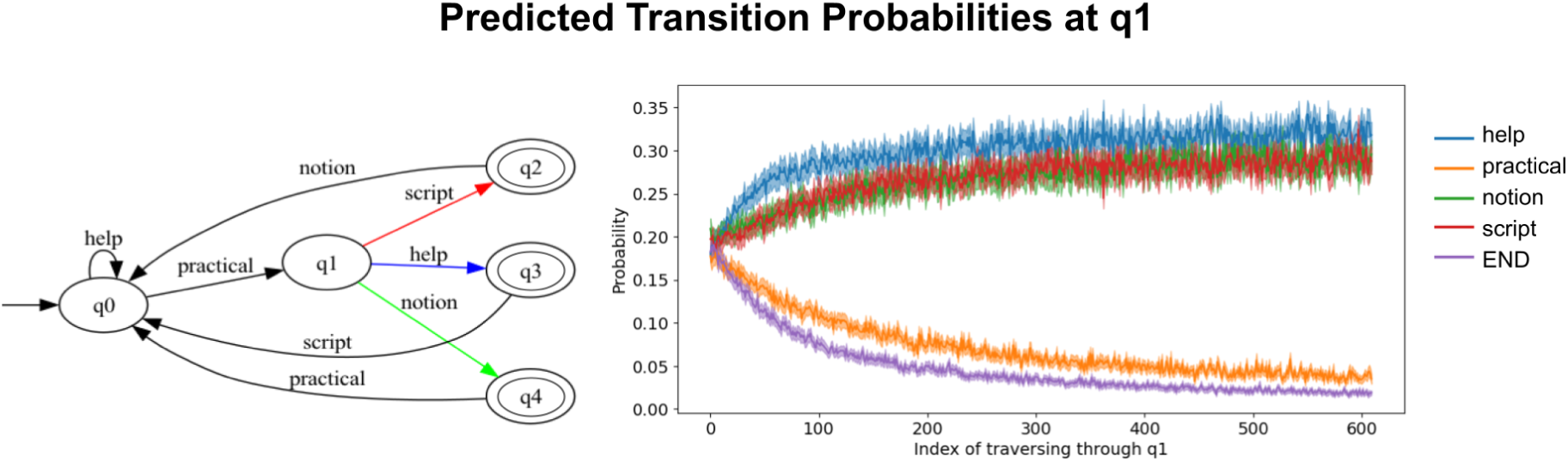
The model predictions for transition probabilities at q1. The true transition probabilities at q0 are 0.33 to script, help, and notion. We indeed see that the model-assigned probabilities converge around the true probabilities. It is also replicated in other DFA’s states (Fig. S8).

We indeed see that the model successfully differentiates between two groups of words: those that are not valid continuations from q0, whose probabilities decrease to zero, and those that are valid continuations from q0, whose probabilities increase and stabilize roughly around 0.3. This finding demonstrates the LLM’s ability to identify multiple valid continuations at each step, supporting the claim that it captures all possible valid sequences in the regular language defined by the underlying structure.

Additionally, there is a noticeable difference in the probabilities assigned to “help” compared to “script” and “notion.” This disparity may be attributed to the higher frequency of “help” in the regular language (half of the valid sequences in the language start with “help”), which could lead to a model bias toward this word.

As a complementary analysis, we compute the KL divergence between the model’s predicted next-token distribution and the true transition probabilities at each state (Fig. 7). Following the transition from the random baseline to the structured phase, KL divergence briefly increases before rapidly converging, indicating alignment between the model’s predictions and the underlying statistical structure.

**Figure 7:**
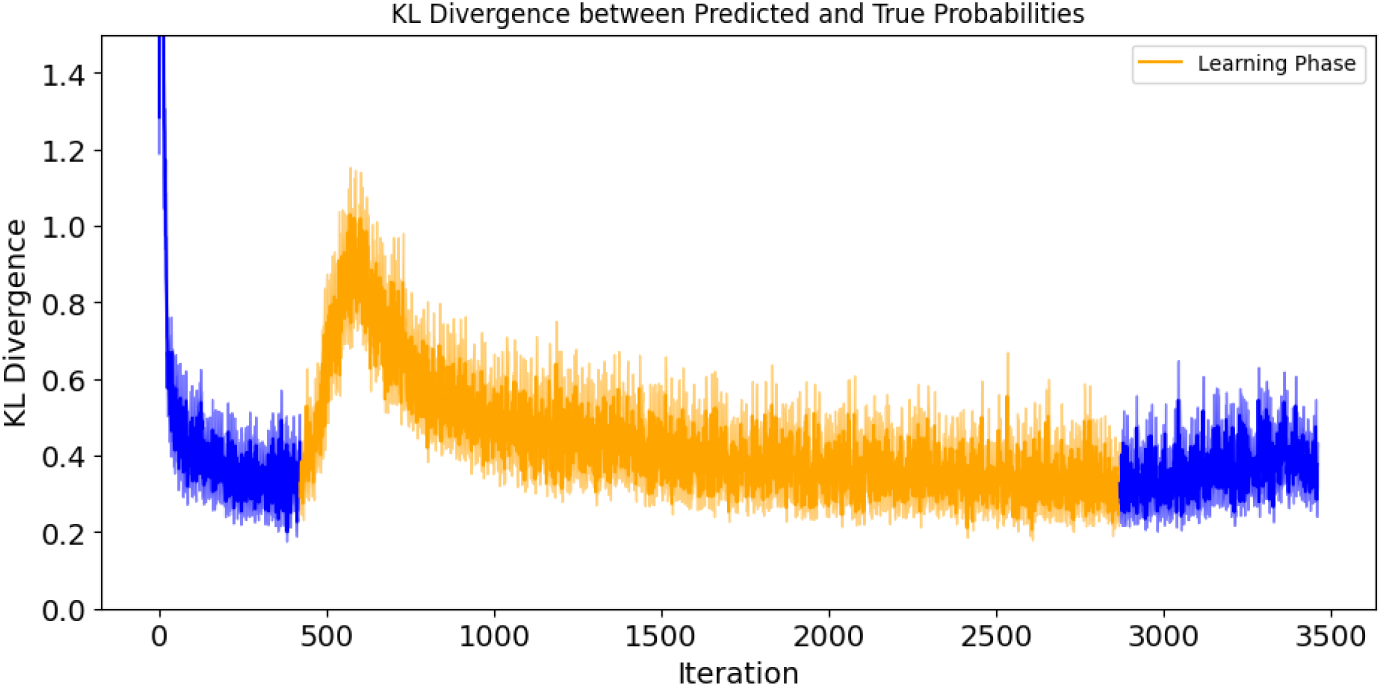
KL divergence between predicted and ground-truth probabilities across training. The orange region marks the learning phase, showing the model’s improved alignment with the Bayes-optimal predictor.

## Discussion

Our findings highlight that large language models (LLMs) can solve statistical learning tasks, which traditionally thought to depend on neural plasticity, through computation alone, without weight adaptation. This result challenges a long-standing assumption in cognitive science that adaptive behavior necessarily requires changes to the system’s structure. Instead, it suggests that sufficiently trained predictive systems may flexibly generalize to novel tasks using their existing representational resources. For the study of human cognition, this opens the possibility that some forms of learning attributed to synaptic plasticity might instead be supported, at least in part, by dynamic computations operating over pre-existing knowledge.

Growing evidence suggests that learning processes occur at the level of individuals, clusters and networks of neurons. At the individual neurons, through intrinsic dynamic activity patterns of bioelectric circuits (Banerjee, 2015; Biswas et al., 2021, 2022; Law & Levin, 2015; Snipas et al., 2016). At the cluster level, recent work in the gustatory cortex has shown that clusters of biological neurons driven by learned anticipatory cues accelerate sensory coding to expected stimuli (Mazzucato et al., 2019). And at the network level, it was demonstrated that rapid motor adaptation can arise from population-level dynamics in the premotor cortex – without any detectable change in synaptic connectivity (Perich et al., 2018). These and others challenge the notion that learning takes place through synaptic (or weight adaptation) alone. However, it is unclear how these can be reconciled with the fact that statistical learning is persistent over long periods (Kóbor et al., 2017)—supporting the existence of additional neurobiological processes.

Despite evidence that statistical learning in humans relies on neural plasticity (Conway, 2020), more recent findings in neuroscience point to other mechanisms that also play a role in learning. Kumaran & McClelland (2012) and A. C. Schapiro et al., (2017) showed models of the hippocampus in which synaptic plasticity is used to learn the training dataset. Still, the underlying regularities are not encoded in the model weights. Then, generalization in this network occurs through recurrent connections without additional training. These models demonstrate how the brain may use computations to solve statistical learning tasks by integrating knowledge to find regularities that were not explicitly encoded in weights. Sorscher et al., (2022) showed a biologically plausible model for the brain in which extensive training yields representations that require very little plasticity to adapt to new labels. This model demonstrates how rich representations can be used for out-of-distribution generalization, which could compensate for the lack of data samples in learning tasks.

While prior work has shown that intrinsic dynamics can give rise to clustering patterns (Kolen, 1994) and that transformer models exhibit characteristic behaviors in embedding space even without training (Singh, 2023), our findings demonstrate that pre-training is essential for the phenomena we observe (Fig. S11). These results suggest that the effects we report cannot be attributed solely to the model’s inherent architecture or dynamics.

Recent research has increasingly converged on the idea that LLMs offer a promising computational framework for understanding human cognition and brain function. Empirical studies have shown that LLM activations correlate with neural responses across multiple regions involved in language processing, suggesting shared computational principles between artificial and biological systems (Caucheteux & King, 2022; Goldstein et al., 2022, 2023, 2024; Schrimpf et al., 2021). This convergence is particularly salient in the context of predictive processing, where both brains and LLMs exhibit forward-looking, context-sensitive behavior (Binz & Schulz, 2023; Tuckute et al., 2023). More recent work has extended these findings, exploring how LLMs mirror hierarchical and semantic structures observed in human cognition (Ren et al., 2024; Yao et al., 2025), and how their internal representations can reconstruct the temporal dynamics of brain activity during language comprehension (Kumar et al., 2022). Moreover, theoretical reviews have begun to clarify both the opportunities and the limitations of using LLMs as cognitive models, noting that while they do not replicate mechanisms of biological learning, their emergent behaviors may nevertheless capture key aspects of human-like reasoning, memory, and abstraction (Binz & Schulz, 2023; Niu et al., 2024; Sartori & Orrù, 2023). Together, these findings point to LLMs not merely as engineering tools, but as valuable model organisms for probing the structure and function of human thought. In conjunction with the findings of our study, this notable similarity offers potentially new insights into generalization mechanisms in the brain, namely that prior knowledge of the learning system can be leveraged to perform new tasks without plasticity. We do not claim that LLMs provide a one-to-one mapping onto cognitive mechanisms. Rather, we argue that there are meaningful similarities between the underlying processes. For instance, while LLMs possess a context window spanning thousands of tokens, far exceeding the capacity of human memory, their *effective* memory (i.e., the ability to retrieve information from prior input) decays with increasing distance from the current token (Liu et al., 2024). This degradation resembles the temporal decay observed in human memory (Oberauer & Lewandowsky, 2008), potentially pointing to shared constraints. However, the nature and extent of this parallel remain uncertain (Vaidya et al., 2023).

This paper questions some of the main assumptions regarding learning and its relation to change in the learning system. First, we suggest that the brain may rely on computations rather than weight adaptation more than previously acknowledged. This follows A. C. Schapiro et al., (2017) findings, which proposed a model for the hippocampus that could support simple statistical learning before having the opportunity to encode the statistical regularities in its weights. The generalization in this network occurred during inference time, using recurrent connections. The idea of achieving generalization via computation was supported from LLMs perspective. Geva et al., (2021) showed that transformer-based LLMs encode memories within their feed-forward layers. Still, the prediction is achieved in inference time via an efficient computation of those encoded memories. Our findings integrate the above ideas into a broader perspective without assuming specific neural mechanisms or cognitive processes.

Second, we show that sufficiently trained predictive systems can generalize to complex regularities apparently far from their training data distribution. To clarify what “sufficiently trained” means, it is important to acknowledge the unpredictability of when new capabilities emerge. Although training scale and data quality are critical ^48^, the shift from basic to advanced abilities is not well defined. Models such as GPT-3 ^3^ and PaLM ^49^ illustrate that larger size and richer datasets often coincide with emergent skills, though the exact thresholds remain uncertain. In this sense, a system may be considered sufficiently trained once its scale and data coverage allow unforeseen abilities to arise. Along similar lines, Sorscher et al., (2022) suggested biologically plausible modeling for the brain, showing that representations similar to those in the inferotemporal (IT) cortex can help to identify new visual concepts using a simple plasticity rule. They suggested that extensive pre-training yields rich representations that enable the applicability of such a simple plasticity rule. Our work shows even more impressive capabilities of a pre-trained learning system, as we show that they could generalize to patterns and structures far from any of their training data distribution without adaptation. One might still question, however, whether such generalization truly reflects abstraction, or instead arises from memorized patterns in the training data. While the exact composition of training data for most LLMs is not publicly available, we believe that the specific sequences used in our experiments are highly unlikely to have been encountered during training. Our sequences are generated from abstract DFAs with randomly assigned vocabularies that lack any semantic content, syntactic structure, or executable logic. As such, they bear no resemblance to the natural language that LLMs are trained on.

Last, our findings suggest a distinction between the computation producing the adapted output and the system’s state change. While learning in the brain is assumed to happen in tandem (Neves et al., 2008; Takeuchi et al., 2014), in-context learning proposes that they can be observed disjointly. I.e., it is possible that the brain can adapt to a new input via computation alone, and this adaptation does not inherently require plasticity. Plasticity is then used to encode the adaptation within the system parameters, for a longer-term process. This raises further questions about the distinction between memory, learning, and generalization, and we hope that this work opens the door for future neuroscience research to use deep learning and AI models to provide novel insights into foundational concepts from the cognitive and neuroscience communities.

## Acknowledgments

This work was supported by the European Research Council (ERC) under the European Union’s Horizon Europe research and innovation programme (ERC Starting Grant “WordOrigions”, Grant Agreement No. 101222215) and by the Israeli Science Foundation grant no. 1023/26. We also gratefully acknowledge our colleagues’ support and contributions to the reviewers’ insightful feedback: Prof. Haim Sompolinksy, Prof. Baruch Eitam, Dr. Sebastian Michelmann, Dr. Noam Siegelman, Dr. Yael Bitterman, Mr. Itay Itzhak, Ms. Daria Lioubashevski.

## Supplementary Materials

### Materials and Methods

#### A Sequence Generation for The Experiments

To generate sequences for our experiment, we employ a random walk on a graph that is designed to reflect the structure of the DFA representing a regular language, with an additional final state incorporated into the design. This process is illustrated in Fig. S1.

The graph structure is similar to the DFA, as each node in the graph corresponds to a state in the DFA, and the edges correspond to the DFA transitions. We add an additional node, in which each node corresponding to an accepting state of the DFA has an outgoing edge to that additional node.

We then perform a random walk on this graph, starting at the node corresponding to q0, and ending at the additional node mentioned above. During each step of the random walk, a transition is made to a neighboring node according to a random choice of the available edges. Each edge in the graph is associated with a specific word, corresponding to the DFA transitions. So, in each step of the random walk, we add to the sequence the word corresponding to the edge that was randomly chosen between all outgoing edges. This randomness introduces variability in the sequences while maintaining the structural constraints of the DFA. The sequences generated by this process inherently conform to the structural rules defined by the DFA. The inclusion of the final state ensures that each sequence is complete and valid.

This approach allows us to create a variety of sequences that adhere to the regular language’s structural constraints, enabling a robust evaluation of the LLM’s ability to learn and predict based on such sequences. The random walk method ensures that our sequences exhibit the complexity and variability necessary for meaningful experimentation while maintaining adherence to the underlying DFA structure.

### A.1 Technical Details

SRTT: The process begins with 80 randomly generated sequences, totaling 496 tokens, with sequence lengths varying randomly. Next, during the structured phase, the model receives 320 structured sequences, contributing 2,880 tokens. Finally, the model is presented with another set of 80 random sequences, totaling 478 tokens. In total, this sequence comprises 480 sequences and 3,954 tokens. For vocabulary coverage, we conduct the experiment using 100 distinct vocabularies, each containing four unique tokens, resulting in a total coverage of 400 unique tokens.

AGL: We start with 85 randomly generated sequences, totaling 422 tokens. During the structured phase, the model is fed 340 structured sequences, totaling 2,449 tokens. Finally, another 85 random sequences are introduced, adding 407 tokens. In total, this sequence consists of 510 sequences and 3,278 tokens. For vocabulary coverage, we conduct the experiment using 100 distinct vocabularies, each containing 4 unique tokens, resulting in a total coverage of 400 unique tokens.

Model: mistralai/Mistral-7B-v0.3

Context window length: 32768

Number of parameters: 7,248,023,552

Hugging Face URL: https://huggingface.co/mistralai/Mistral-7B-v0.3

Model: meta-llama/Llama-3.1-8B

Context window length: 131072

Number of parameters: 8,030,261,248

Hugging Face URL: https://huggingface.co/meta-llama/Llama-3.1-8B

## B Calculating the theoretical upper bound for transition probabilities

The DFA representing the task induces transition probabilities for the graph traversing. In this paper, we compare how close are the probabilities that the LLM assigns to the transition probabilities that the DFA induces, referring to it as the theoretical upper bound. Let S be a sequence induced by traversing the following DFA states:

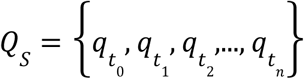

Then, the theoretical upper bound for the sequence is the average of all transition probabilities between each pair of consecutive states:

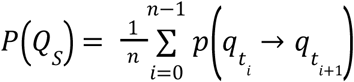

In this paper, the theoretical upper bound is calculated over the complete input sequence, which is 1 for the SRTT, as each state has only one outcoming edge, and for the AGL is 0.45, as most states have two outcoming edges (with probability 0.5), and one state has three outcoming edges (with probability 0.33).

## C Probabilities over sequences

In the paper, we presented the probability over each word, and below, we accumulated the word probabilities into sequence probability. Each sequence is separated by a special ‘END’ token, which represents traversing from an accepting node back to q0, such that between each two ‘END’ tokens, there is a valid sequence that belongs to the vocabulary of the DFA.

### C.1 Mean on sequence

First, we calculate the probability of a sequence as the mean of all predicted probabilities of its words, given all previous tokens:

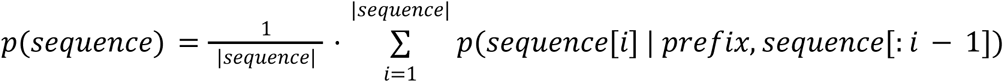

For example, if we want to calculate the sequence probability of “practical notion” from Fig 2B (iterations 1506-1608):

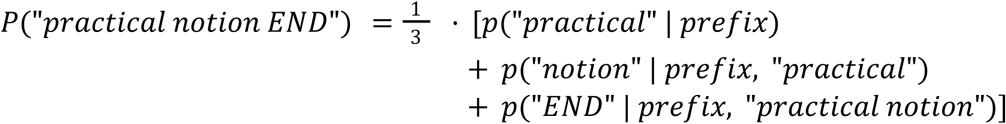

Where *prefix* is the concatenation of the 1505 input words until this sequence?

In Fig. S2a, we see that the model also shows learning at the structured part when calculating the mean of a sequence.

### C.2 Multiply the probability of the tokens combining the sequence

In LM evaluation, it is common to factorize the joint probability over a sequence as the product of conditional probabilities of the words and symbols combining this sequence (Bengio et al., 2003; Radford et al., 2019).

E.g., we calculate the probability of a sequence as follows:

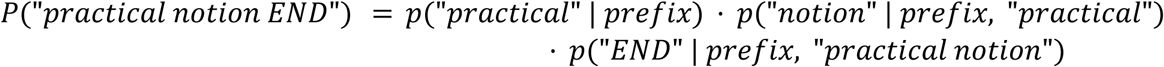

In Fig. S2b, we see the learning in the structured part, as well as in calculating the production of a sequence.

---

**Figure S1:**
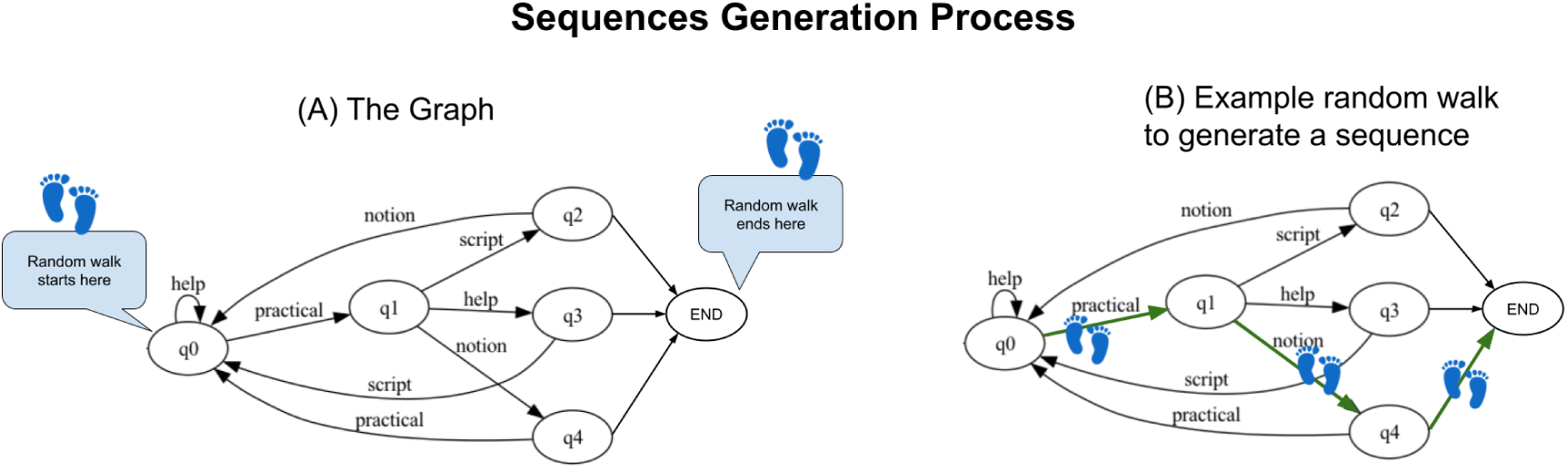
Sequence Generation Process. A illustrates the graph used for sequence generation. A single random walk from the initial state (q0) to the terminal state (END) produces a sequence. B provides an example of such a walk. At each node, an outgoing edge is selected uniformly. For instance, the edge leading to ‘practical’ is chosen with a probability of 0.5, followed by ‘notion’ with a probability of 0.33. The walk terminates at END with a probability of 0.5,resulting in the sequence “practical help END”.

**Figure S2:**
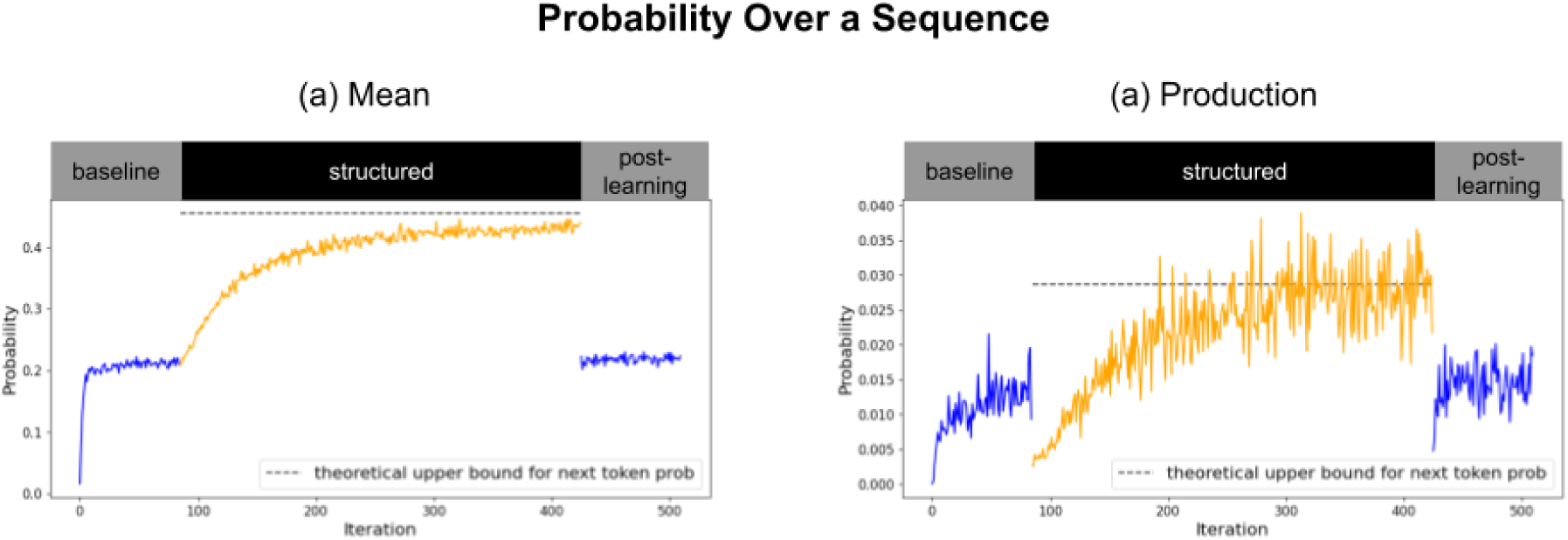
(a) predicted probabilities of a sequence via mean on tokens. (b) predicted probabilities of a sequence via the production of conditioned probabilities of its tokens.

**Figure S3:**
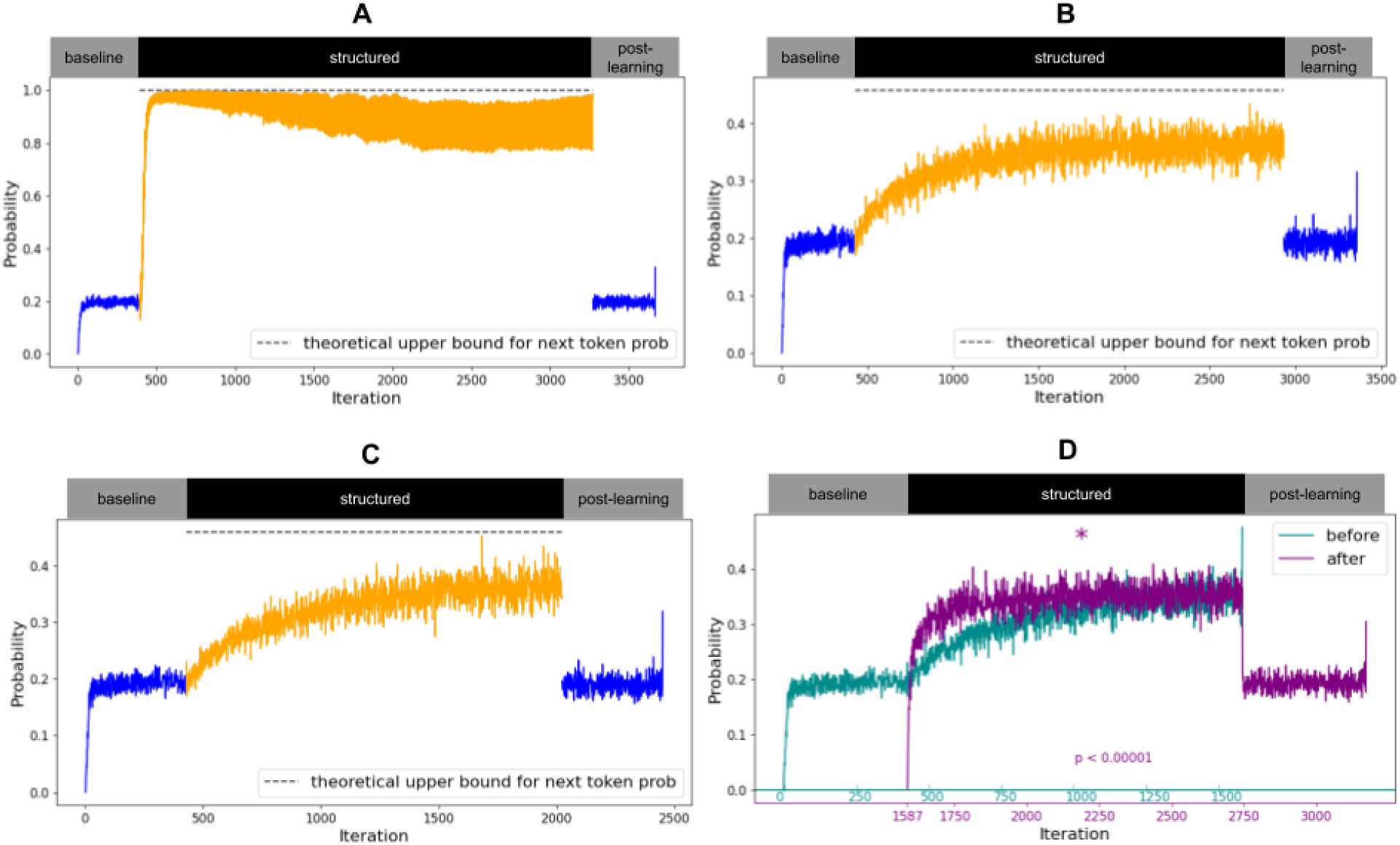
Results of Llama-3.1-8B language model (Grattafiori et al., 2024). (A) SRTT. (B) AGL base setup. (C) AGL switched vocabulary in the middle of the trial. (D) AGL has no duplicates.

**Figure S4:**
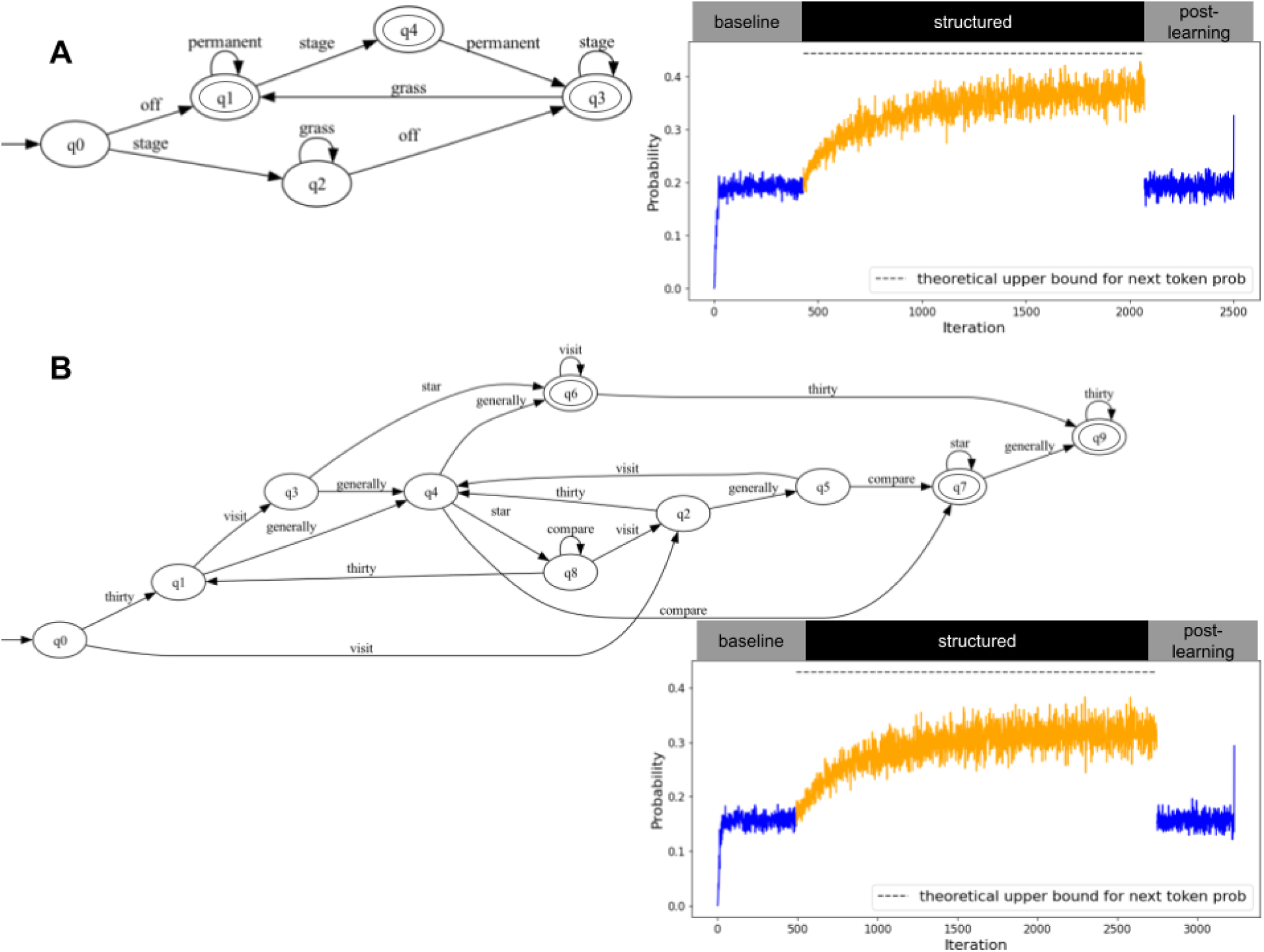
(A) Results of Llama-3.1-8B (Grattafiori et al., 2024) over DFA inspired by Knowlton & Squire (Knowlton & Squire, 1994). (B) results of Llama 2 over DFA inspired by Vokey & Brooks (Vokey & Brooks, 1992).

**Figure S5:**
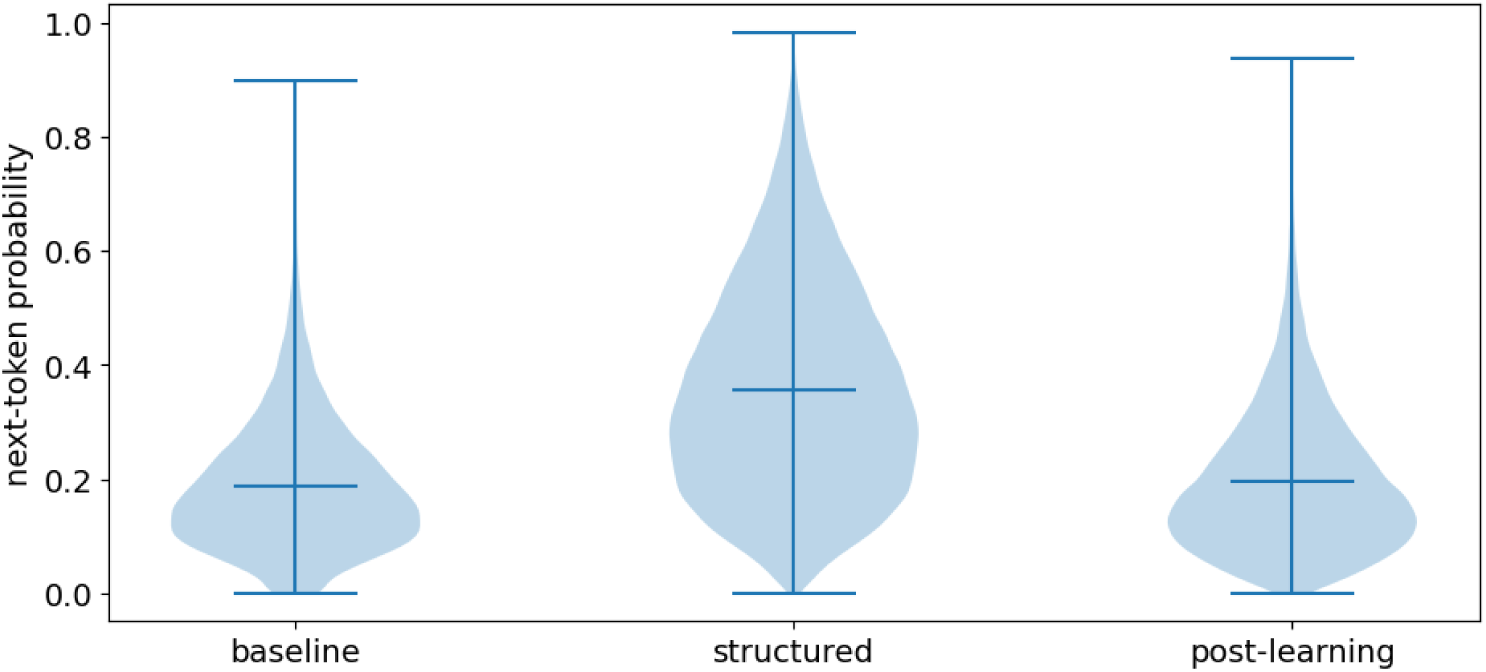
The violin plot illustrates the distribution of predicted probabilities for each part of the experiment. We find a significant difference between structured and random parts, supported by a permutation t-test (t=135.43, p=10.0e-05).

**Figure S6:**
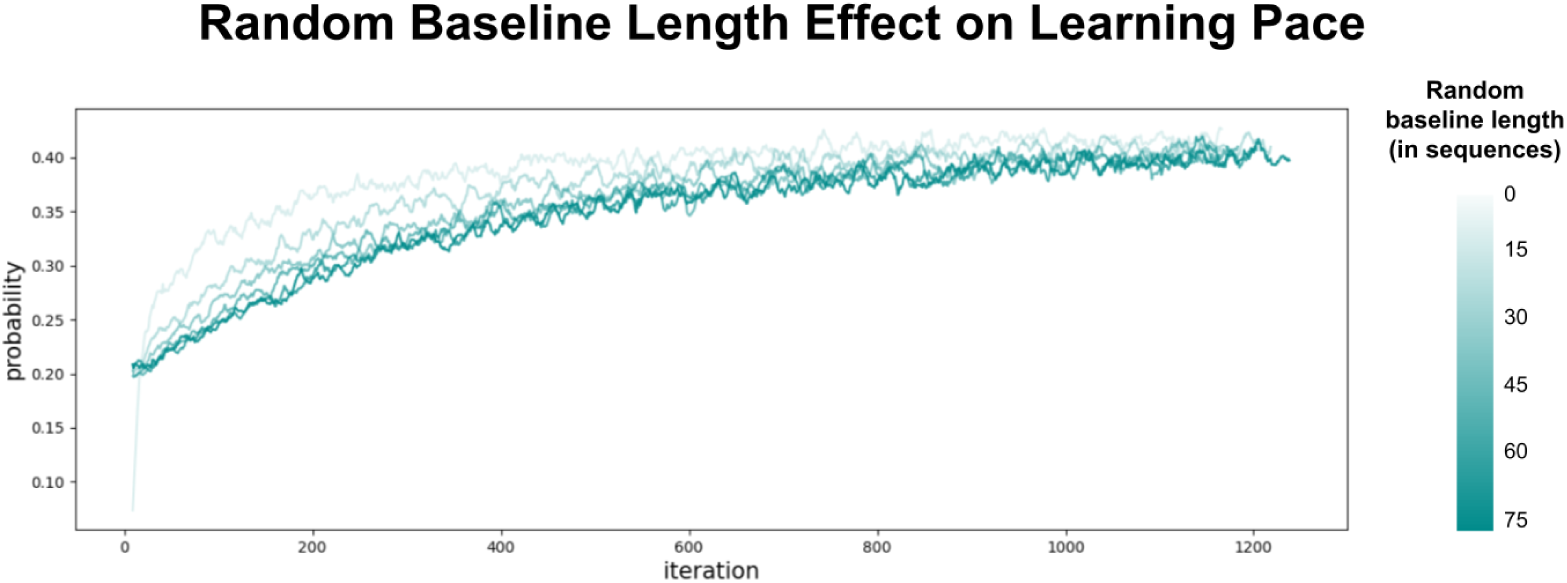
Learning curve for different baseline part lengths (smooth with window size=10). Light represents a shorter random baseline length before the structured part, which shows faster learning, compared to darker green, which represents a longer random baseline at the beginning.

**Figure S7:**
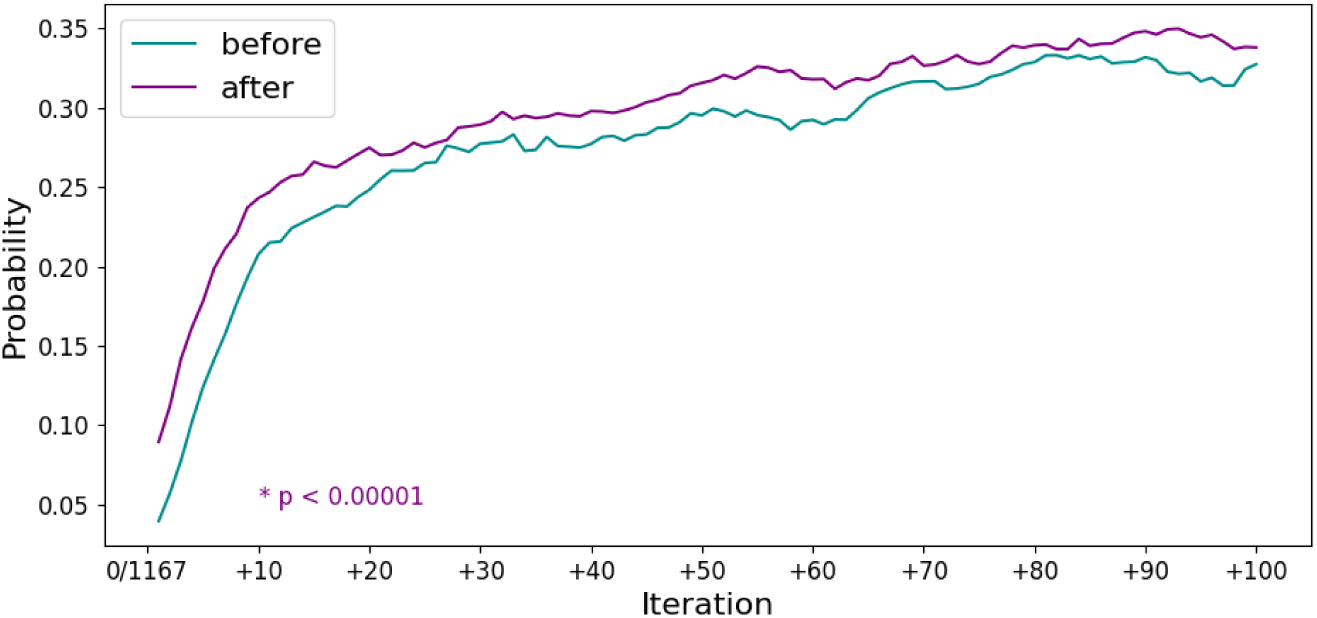
zoom-in over the first 100 iterations (smooth with window size=7) in a setup without any noise at the beginning, i.e., skipping the baseline part. We find that the probabilities after the switch are significantly higher than at the beginning of the structured part, even without any noise.

**Figure S8:**
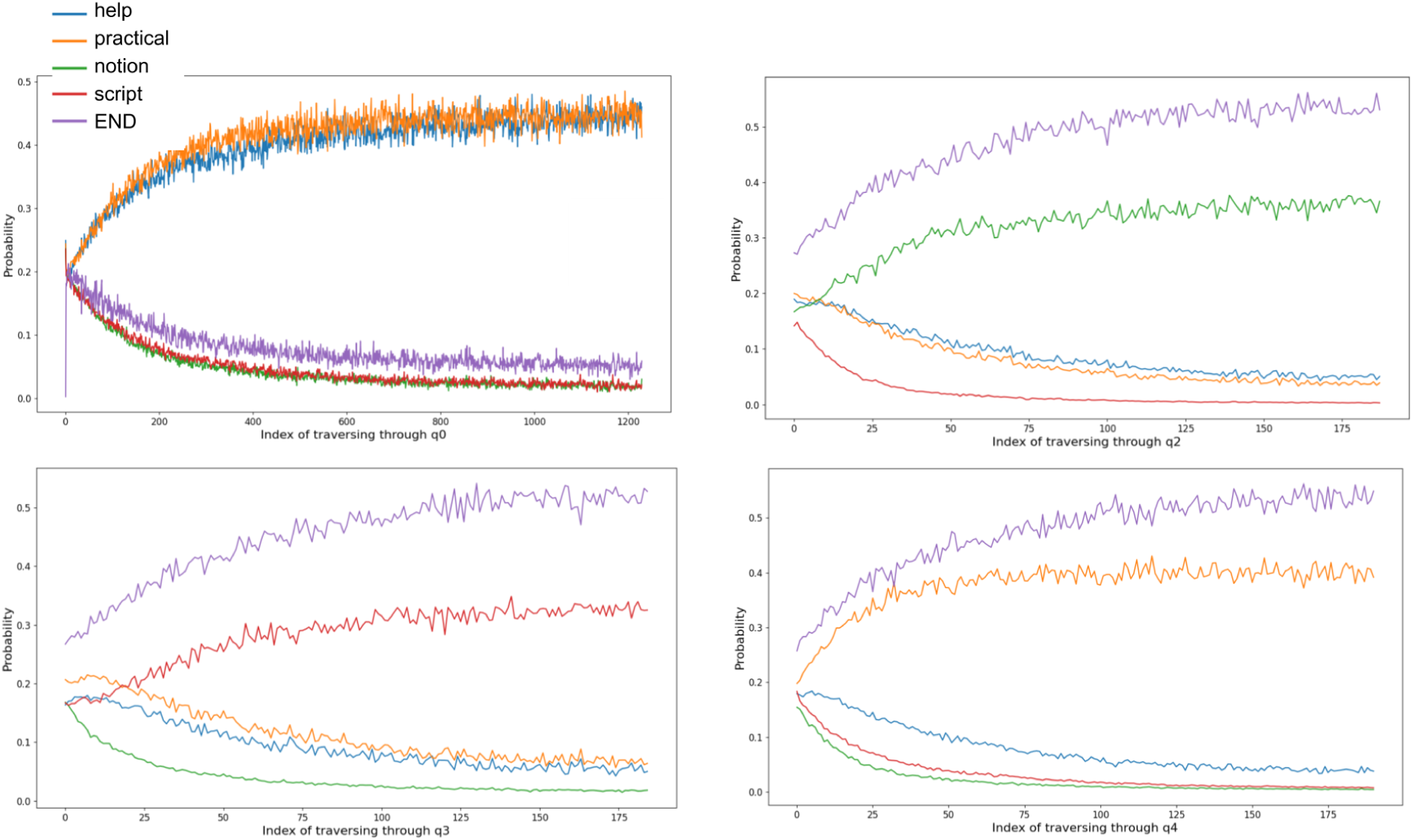
The model predictions for transition probabilities at each stage of the DFA, in addition to q1, are presented in Fig 4. The true transition probabilities are 0.5 for two different words at each state, as each state has exactly two outcoming edges with different labels. We see that, indeed, in all states, the model captured the two words with higher probabilities than the other words that are not the edge labels.

**Figure S9:**
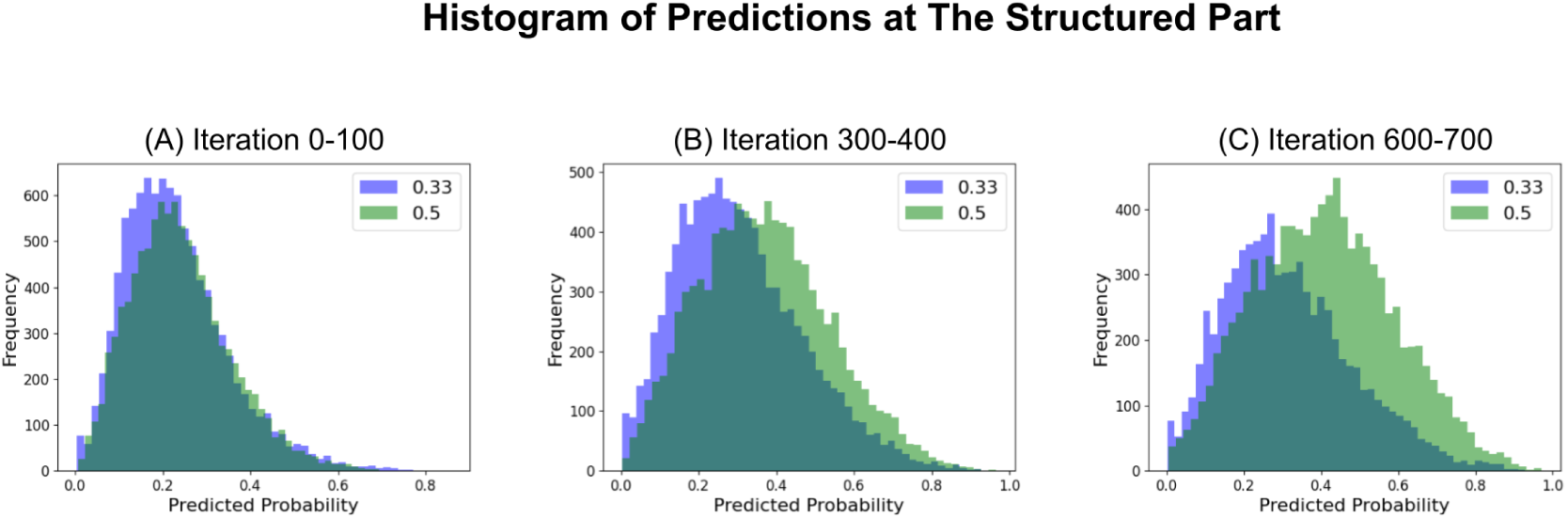
Evolution of Prediction Probabilities Across Experiment Lengths. Histograms of prediction probabilities for all samples, categorized by true probability (0.33 and 0.5). The three subfigures illustrate the model’s learning progression as it encounters longer sequences. Initially (A), predictions for both true probability classes are dispersed, indicating random-like behavior. As sequence length increases (B and C), the distributions become more distinct and centered around their respective true probabilities, demonstrating improved model performance.

**Figure S10:**
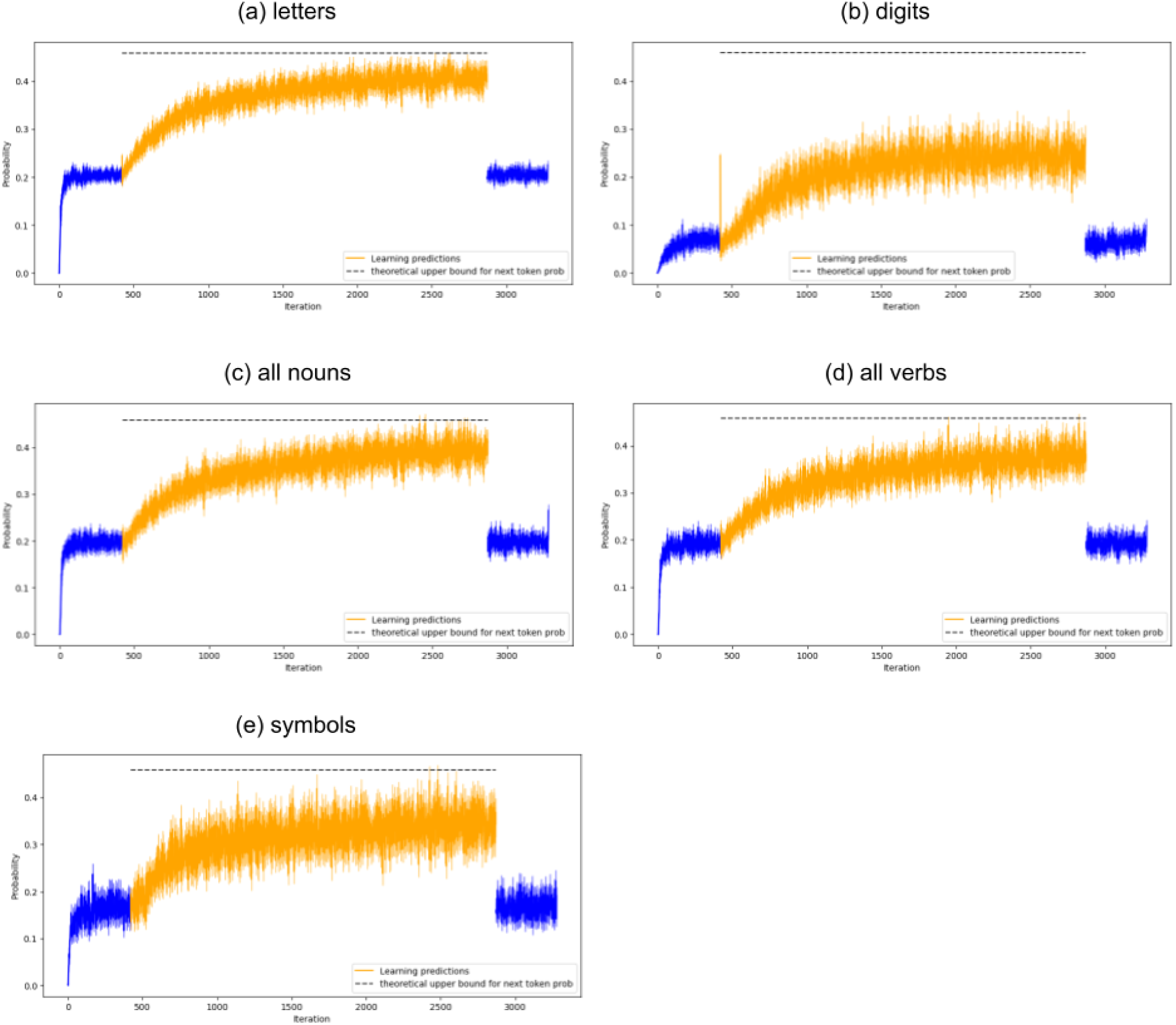
AGL experiment with different types of vocabularies. (a) vocabulary is all letters. (b) vocabulary is all digits. (c) vocabulary is all noun words. (d) vocabulary is all verbs. (e) vocabulary is symbols !@#$%^&*()_+-={}[]±§/\”\’:;.?>,<∼.

**Figure S11:**
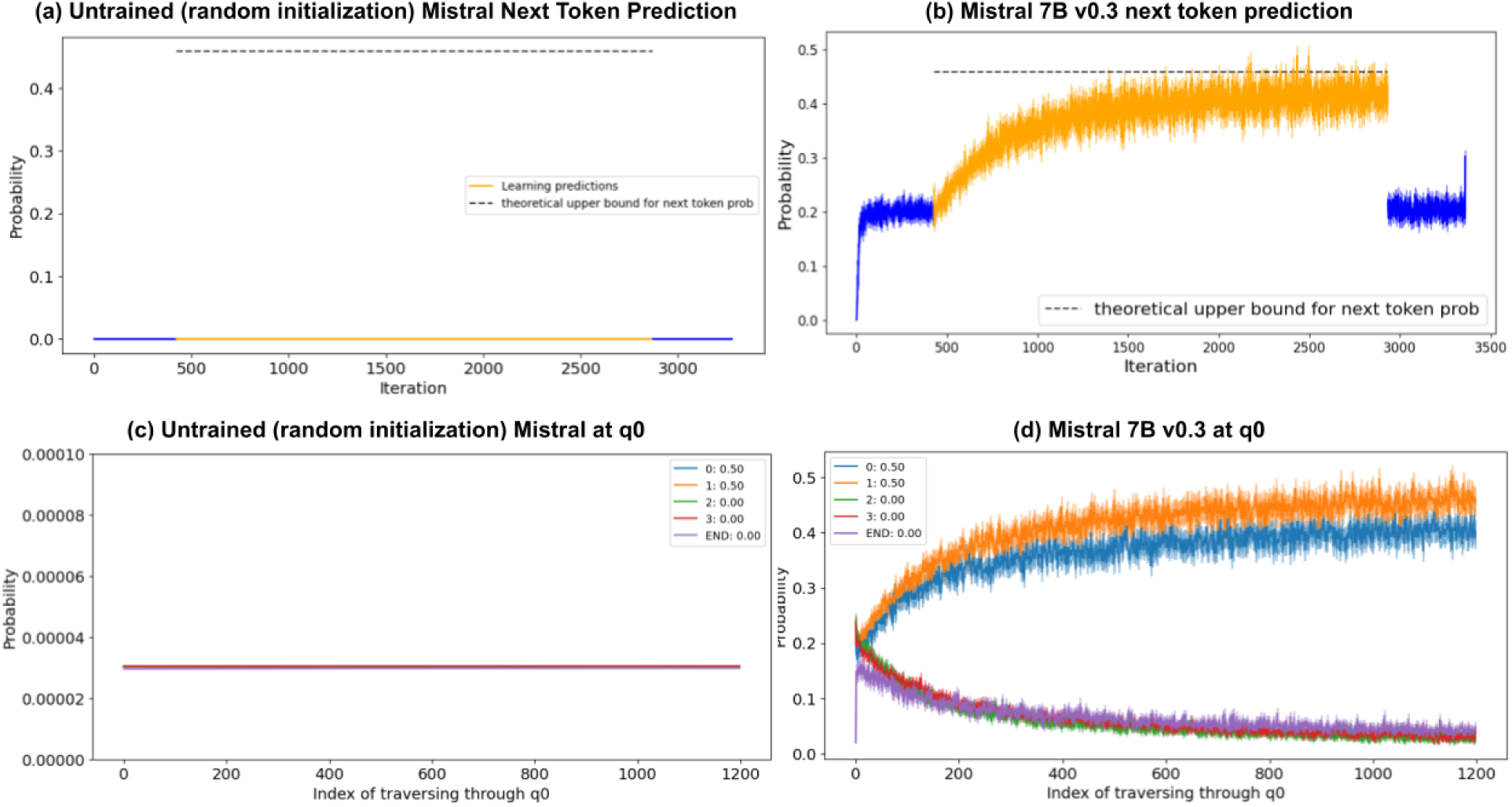
Pre-trained vs. Untrained Network. (a) Next word prediction when running an untrained Mistral model, with randomly initialized weights. (b) Next word prediction for the pretrained model (same as Fig. 3b). (c) Probabilities assigned by an untrained Mistral. (d) Probabilities assigned by the pre-trained Mistral 7B v0.3 model (same as in Fig. 3b). In (c), all tokens in the vocabulary receive nearly uniform probabilities close to 0, whereas in (d), the model differentiates between valid continuations (blue and orange) and invalid ones, assigning significant probability mass to plausible completions. This demonstrates that the ability to capture statistical structure does not emerge purely from the network architecture but is instead acquired through pretraining.

**Figure S12:**
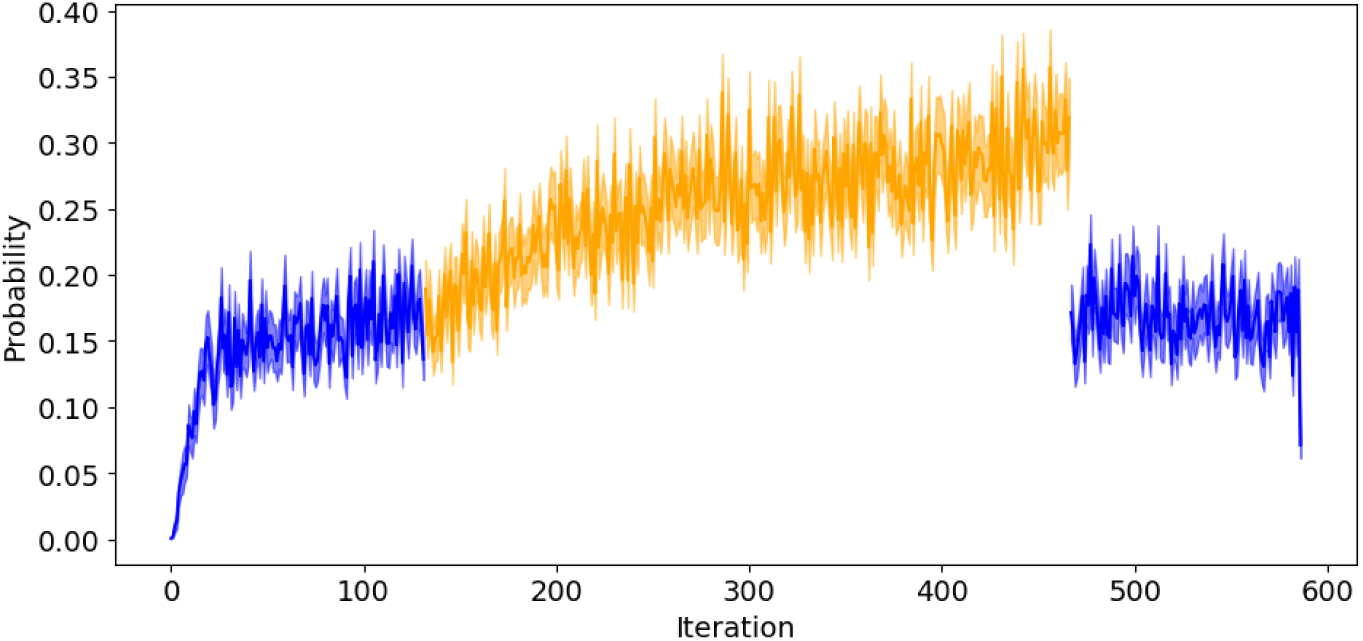
AGL experiment for GPT2-xl (Radford et al., 2019). Replicating the results for the AGL experiment.

## Footnotes

1 Our experiments can be reproduced via our provided <u>Colab notebook</u>.

2 We specifically use the Mistral-7B-v0.3, with a context window size of 32k.

